# Fmp30 is a phosphatidylinositol hydrolase that regulates CoQ biosynthesis

**DOI:** 10.1101/2025.05.01.651778

**Authors:** Zakery N. Baker, Rachel M. Guerra, Sean W. Rogers, David J. Pagliarini

## Abstract

Coenzyme Q (CoQ, ubiquinone) is a redox-active isoprene lipid that supports fundamental enzymatic and antioxidant roles in mitochondria and beyond. Despite CoQ’s importance in organismal health and disease, the mechanisms that regulate its biosynthetic pathway remain elusive. Motivated by this, we mined *S. cerevisiae* multiomics datasets for genes whose disruptions alter CoQ levels and discovered the gene *FMP30* as an unexpected negative regulator of CoQ biosynthesis. Loss of *FMP30* results in elevated complex Q protein abundance, increased number and intensity of CoQ domains, and ultimately enhanced CoQ biosynthetic flux. We demonstrate that Fmp30, a member of the metallo-β-lactamase super family, displays phospholipase type D activity toward phosphatidylinositol and phosphoinositides, and that its deletion causes marked elevation of these lipid species in purified mitochondria. Collectively, our work nominates Fmp30 as a novel regulator linking mitochondrial phospholipid metabolism to CoQ biosynthesis.

## INTRODUCTION

Coenzyme Q (CoQ) is an essential redox-active lipid responsible for transferring electrons in the mitochondrial electron transport chain, acting as a key cofactor in biosynthetic processes, and preventing oxidative damage throughout the cell^1^. Nearly all organisms synthesize CoQ endogenously, with eukaryotic production occurring at the inner mitochondrial membrane^2^. Defects in CoQ biosynthesis result in diverse human pathologies, including ataxias and nephropathies^3^, and secondary CoQ deficiency is a common feature of numerous mitochondrial disorders^4^. Despite that, limited therapeutic options exist to ameliorate CoQ deficiencies, as exogenous supplementation is hampered by the extreme hydrophobicity of CoQ^5^. To this end, elucidating mechanisms that regulate CoQ biosynthesis may motivate alternative therapeutic interventions that bolster endogenous CoQ production.

The critical nature of CoQ in cellular functions suggests that its synthesis and turnover may be subject to multiple levels of regulation, as is true for other bioactive lipids^6,7^. Early work linked the regulation of CoQ biosynthesis to lipid metabolism and oxidative stress responses via PPARα^8^ and NK-κB^9^, though the mechanisms remain unexplored and appear to be context- and species-specific. More recent studies have hinted at coordinated regulation of CoQ biosynthesis with other important mitochondrial processes, such as complex I assembly^10^, mitochondrial biogenesis^11^, and mitochondrial protein processing and import^12,13^. Moreover, a regulatory relationship between mitochondrial phospholipid metabolism and CoQ biosynthesis has been postulated^14^. However, we still lack a detailed understanding of how cells sense and integrate environmental or intracellular stimuli to modulate their CoQ content. As such, the CoQ regulatory landscape remains a ripe area for discovery with potential therapeutic benefit.

In this study, we undertook a CoQ-focused reanalysis of our existing multiomics dataset^15^ and revealed a surprising link between CoQ abundance and the *S. cerevisiae* gene *FMP30*. We characterize Fmp30 (<u>F</u>ound in <u>m</u>itochondrial <u>p</u>roteome 30; Ypl103c), a mitochondrial intermembrane space-localized protein belonging to the metallo-β-lactamase superfamily, as a lipid hydrolase with specificity toward phosphatidylinositol and phosphoinositides. Collectively, our data establish Fmp30 as a novel regulator connecting mitochondrial membrane phospholipid composition to CoQ biosynthesis.

## RESULTS

### *FMP30* deletion increases the CoQ to PPHB ratio

Coenzyme Q (CoQ) biosynthesis involves the import of cytosolic precursors into the mitochondrial matrix, the attachment of the headgroup to the polyisoprenoid tail, and the subsequent chemical headgroup modifications to form mature CoQ (Fig 1a). To find potential regulators of this process, we examined our Y3K dataset—a collection of multiomic measurements from yeast deletion strains lacking sentinel or uncharacterized mitochondrial genes^15^—to identify yeast knockouts with elevated CoQ content (Supp Fig 1a). Intriguingly, many strains with significantly increased CoQ levels lacked genes related to mitochondrial phospholipid metabolism (Supp Fig 1b). Given the precedent of specific mitochondrial membrane phospholipids modulating CoQ biosynthesis protein functionality^16,17^, we opted to pursue these strains further. To confirm and extend these observations, we generated new deletions of the top phospholipid-related genes from this Y3K analysis, along with select genes representing each major mitochondrial phospholipid biosynthetic pathway (Supp Fig 1c), in a BY4742 background and measured levels of both CoQ and its biosynthetic precursor, polyprenyl-hydroxybenzoate (PPHB). Recapitulating the Y3K dataset, we found a significant CoQ increase in a subset of these strains, matched by a corresponding decrease in their PPHB levels (Fig 1b, c). In these affected strains, the ratio of CoQ to PPHB was >20 fold (Fig 1d), suggesting enhanced CoQ biosynthesis efficiency.

**Figure 1.**
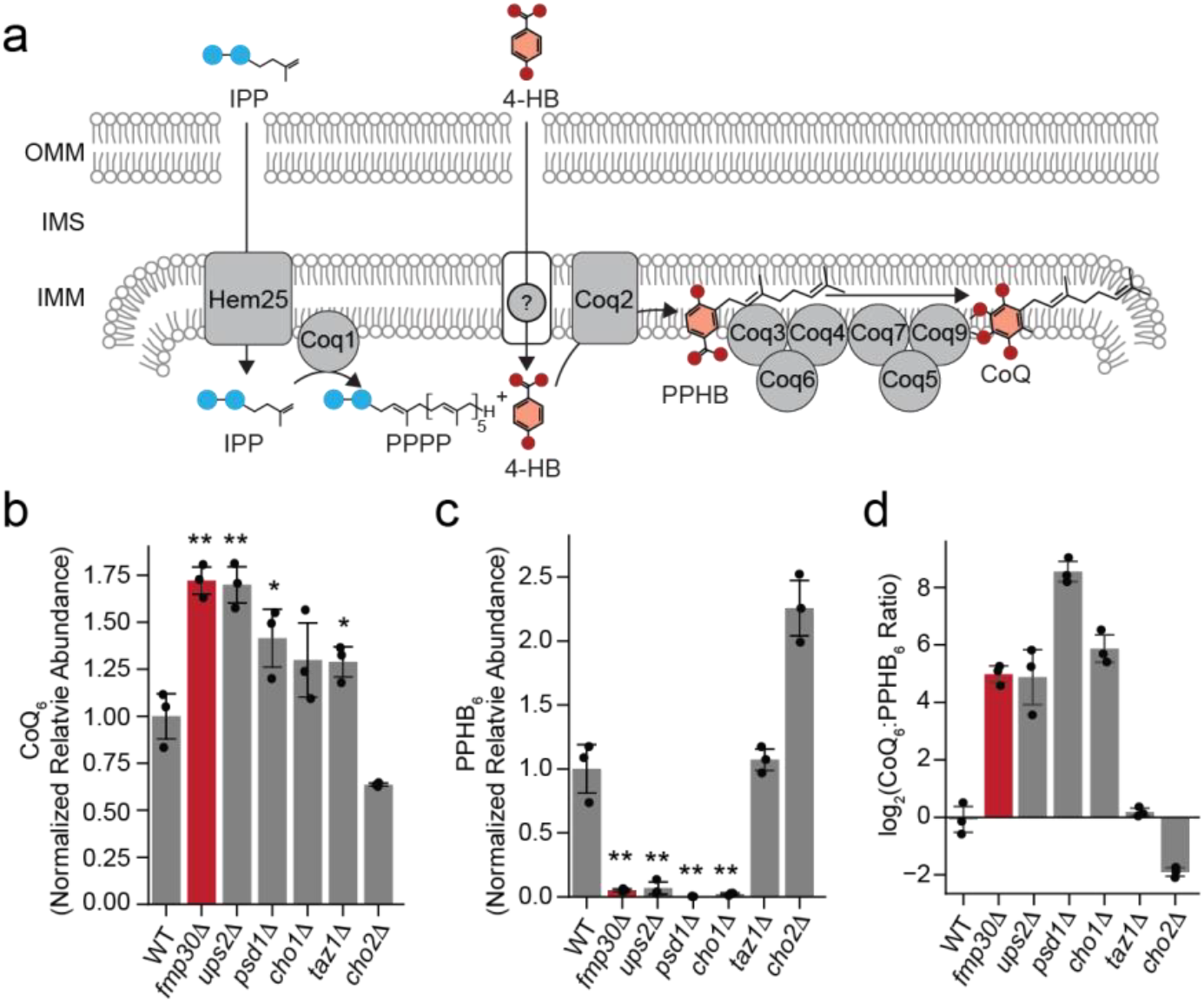
*FMP30* deletion increases the CoQ to PPHB ratio. **a**, Schematic of CoQ biosynthesis in *S. cerevisiae*. Blue circles represent phosphate groups, red circles represent oxygen. IPP, isopentyl pyrophosphate; 4-HB, 4-hydroxybenzoate; PPPP, polyprenyl pyrophosphate; PPHB, polyprenyl-hydroxybenzoate; CoQ, coenzyme Q. **b**, Relative CoQ_6_ abundance in *gene*Δ strains compared to wild type (WT). **c**, Relative PPHB_6_ abundance in *gene*Δ strains compared to WT. **d**, Log_2_ transformed ratio of CoQ_6_ to PPHB_6_ abundance in the *gene*Δ strains in **b** and **c**. *fmp30*Δ is highlighted in red. For **b**-**d**, data are displayed as mean ± s.d., n=3 biologically independent samples, two-sided Student’s *t*-test compared to WT samples. \**P*<0.05, **P*<*0.01.

We set out to determine the underlying mechanism causing increased CoQ biosynthesis focusing on the poorly characterized gene, *FMP30*. Unlike the other affected strains, *FMP30* deletion did not disrupt respiratory growth on non-fermentable carbon sources (Supp Fig 2a), suggesting that our further analyses would not be confounded by global mitochondrial defects. Although *FMP30* has been linked to PE biosynthesis genetically^18^, our untargeted lipidomic analyses of *fmp30*Δ revealed no discernable changes in either whole cell or mitochondrial PE abundance (Supp Fig 2b, Supp Fig 4c). Additionally, when supplemented with ethanolamine at amounts known to boost mitochondrial PE^19^, *fmp30*Δ CoQ levels were not rescued to WT levels. Collectively, this suggests that *FMP30* may have an unappreciated role in regulating CoQ abundance.

### *FMP30* deletion increases CoQ biosynthetic machinery and *de novo* biosynthesis rates

We next measured CoQ biosynthesis rates via pulse-chase labeling of CoQ with ^13^C-labelled 4-hydroxybenzoic acid^16^ . Deletion of *FMP30* increased *de novo* CoQ production at every time point, confirming an increase in biosynthetic efficiency (Fig 2a). To determine the rates of PPHB biosynthesis, we first deleted *COQ7*, which thwarts CoQ biosynthesis beyond PPHB by disrupting complex Q—a metabolon comprised of CoQ biosynthetic enzymes ^15^. *FMP30* deletion in this background did not change the rates of C^13^-labelled PPHB generation, suggesting that the increase in CoQ biosynthesis in *fmp30*Δ occurs at the headgroup modification stage (Fig 1a, 2b).

**Figure 2.**
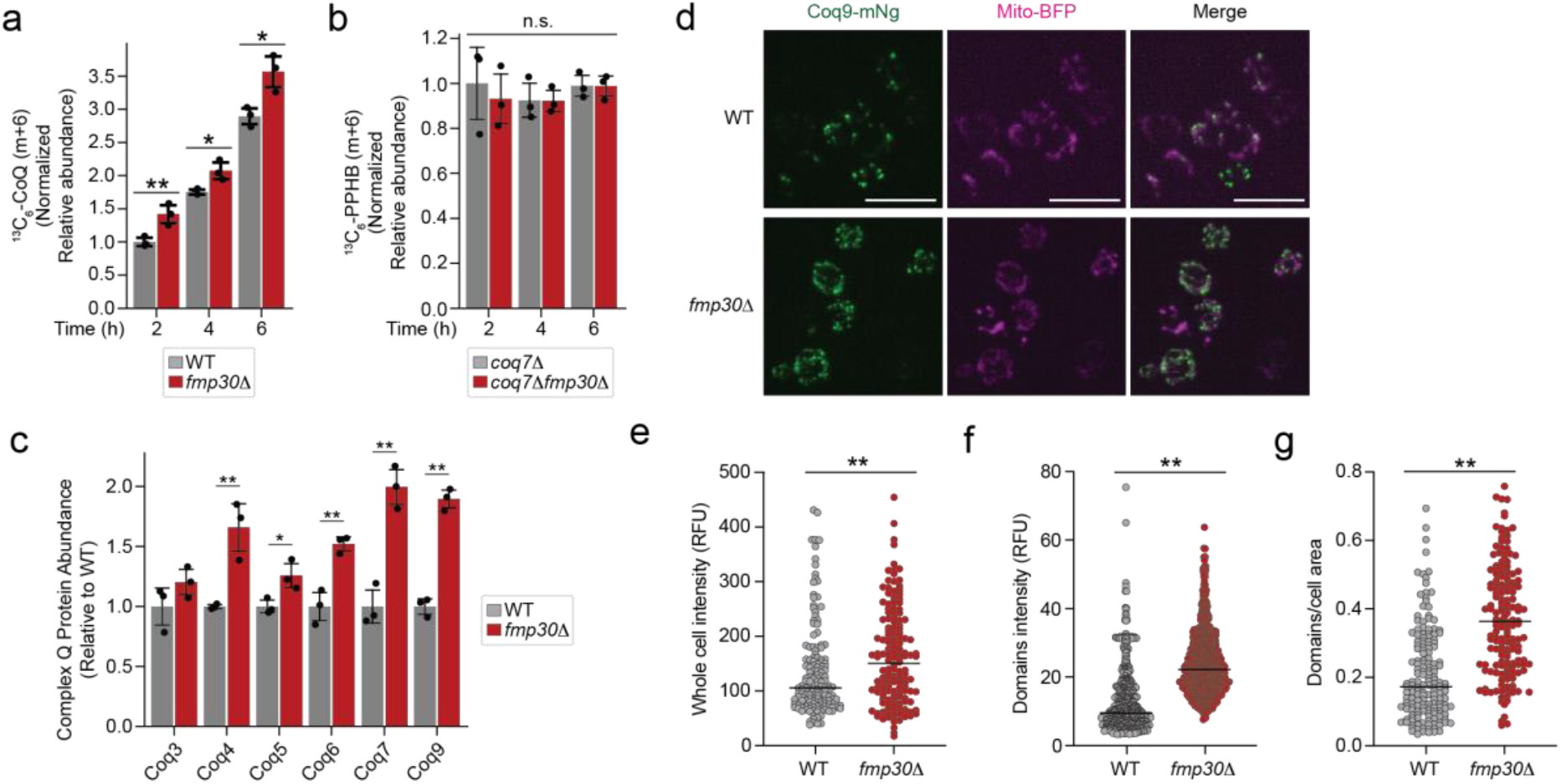
*FMP30* deletion increases CoQ biosynthetic machinery and *de novo* biosynthesis rates. **a**, Abundance of *de novo* synthesized ^13^C_6_-CoQ_6_ in WT and *fmp30*Δ cells grown in -pABA SD 3% (wt/vol) glycerol and 50 μM [*phenyl*-^13^C_6_]-4-HB (relative to WT at 2 hr). **b**, Abundance of *de novo* synthesized ^13^C_6_-PPHB_6_ in *coq7*Δ and *coq7*Δ*fmp30*Δ cells grown in -pABA SD 3% (wt/vol) glycerol and 50 μM [*phenyl*-^13^C_6_]-4-HB (relative to *coq7*Δ at 2 hr). **c**, Relative protein abundance of complex Q proteins from *fmp30*Δ compared to WT. **d**, Representative z-stack images of WT or *fmp30*Δ cells expressing endogenously-tagged Coq9-mNg (green) or Mito-BFP (magenta). Scale bar = 10 μm. (**e**-**g**) Quantification of CoQ domain intensity per whole cell (**e**), individual domain intensity (**f**), or number of domains per cell (**g**) of the cells imaged in **d**. For **a**-**c**, data are displayed as mean ± s.d., n=3 biologically independent samples, two-sided Student’s *t*-test. For **e-g**, all data points are displayed with the median indicated, n>100 cells from three independent experiments, Kolmogorov-Smirnov test. \**P*<0.05, **P*<*0.01.

Complex Q (aka the CoQ synthome), resides on the matrix face of the mitochondrial inner membrane^20,21^. We postulated that elevated complex Q protein stabiliy could boost biosynthetic efficiency. The *fmp30*Δ strain exhibited a slight but statistically significant increase in nearly all complex Q members (Fig 2c), with no global changes to mitochondrial protein content (Supp Fig 2d). Among all strains in the Y3K dataset, *fmp30*Δ possessed the second highest fold change in complex Q proteins (Supp Fig 2d). Beyond abundance, we further assessed how *FMP30* deletion affected complex Q domains—large foci of CoQ proteins that reside near ER contact sites^22,23^ . Indeed, *FMP30* deletion also increased the number and intensity of CoQ domains (Fig 2d-g). Mitochondrial crosslinking coupled with affinity enrichment mass spectrometry of strains expressing tagged Coq9 demonstrate the composition of complex Q remains unchanged in the *fmp30*Δ despite this increased complex abundance (Supp Fig 2e, f). Collectively, these data suggest that increased complex Q and CoQ domain formation in *fmp30*Δ lead to enhanced biosynthetic flux and thus elevated CoQ abundance.

### Fmp30 and human NAPE-PLD lack conserved function

Fmp30 exhibits high overall sequence conservation with the human phospholipase D enzyme, NAPE-PLD (33% identity, 51% similarity) (Fig 3a). Both proteins belong to the metallo-β-lactamase superfamily, sharing the highly conserved HxHxDH motif, which contains the metal-binding residues critical for catalysis (Fig 3b)^24^. We purified both NAPE-PLD and Fmp30, verified their phosphodiesterase activity on a generic substrate Bis-pNPP (Supp Fig 3a), and tested if they could hydrolyze N-acylphosphatidylethanolamines, the native substrate of NAPE-PLD (Supp Fig 3b). While NAPE-PLD rapidly depletes levels of 18:1-PE-N-18:1 (18:1-NAPE) and subsequently generates reciprocal amounts of 18:1 N-acylethanolamide (18:1 NAE), recombinant Fmp30 had no measurable activity toward 18:1-NAPE (Fig 3,d).

**Figure 3.**
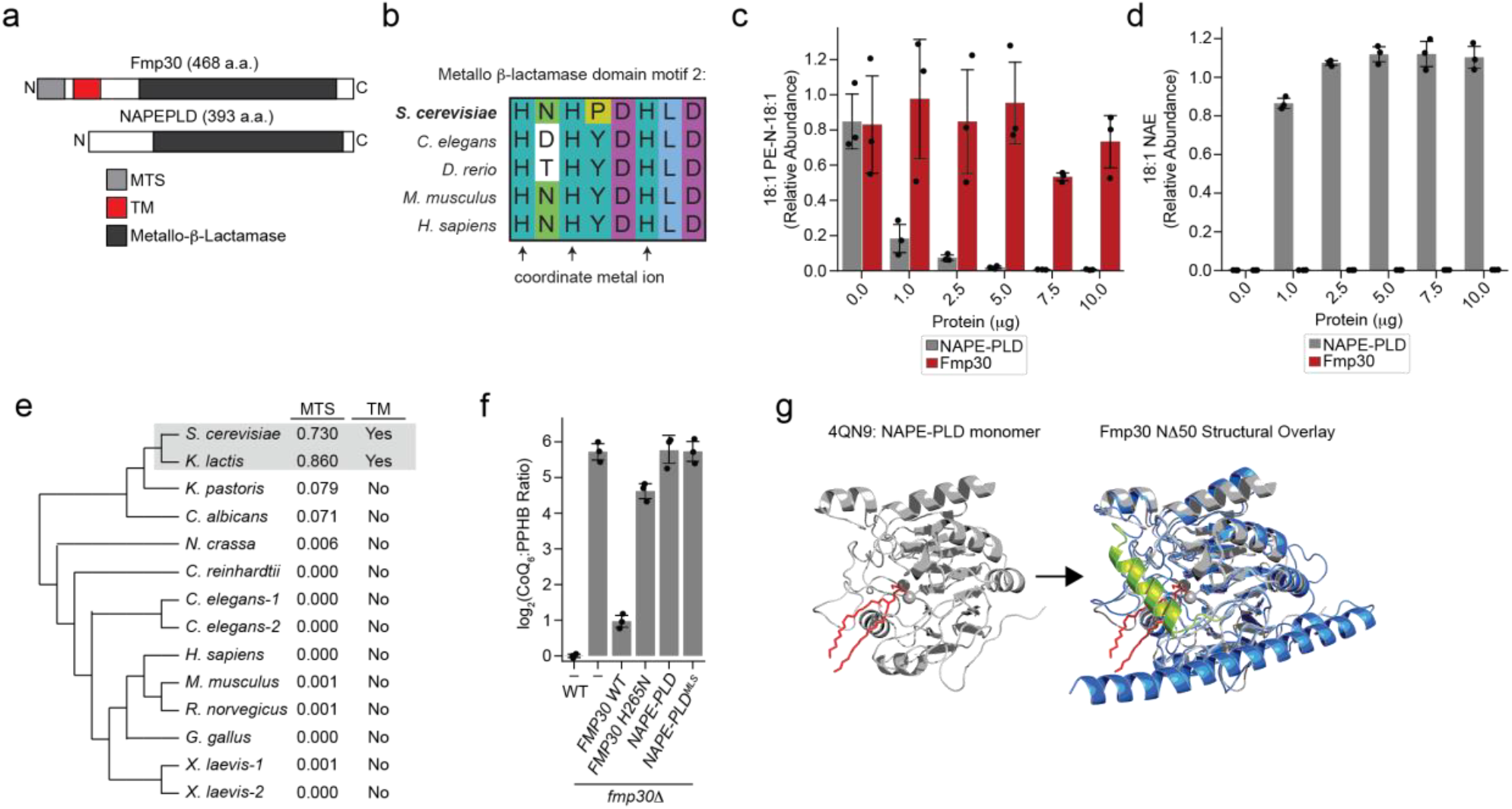
Fmp30 and human NAPE-PLD lack conserved function. **a**, Diagram of Fmp30 and NAPE-PLD protein sequences highlighting protein domains. MTS, mitochondrial targeting sequence; TM, transmembrane domain. **b**, Multiple sequence alignment for metallo-β-lactamase motif 2, highlighting the metal binding site, in NAPE-PLD homologs from various species. Colors represent default ClustalX amino acid class color scheme. **c**, Relative abundance of 18:1 PE-N-18:1 (18:1 NAPE) following incubation with the indicated concentration of purified recombinant Fmp30 or NAPE-PLD for 30 min. NAPE, N-acylphosphatidylethanolamine. **d**, Relative abundance of 18:1 NAE generated after incubation of 18:1 PE-N-18:1 with the indicated concentration of purified recombinant Fmp30 or NAPE-PLD for 30 min. NAE, N-acylethanolamide. **e**, Phylogenetic analysis of Fmp30 homologs. Sequences were analyzed using MitoFates to determine mitochondrial targeting sequence (MTS) probability and AlphaFold structures were analyzed for presence of a transmembrane (TM) domain. **f**, Log^2^ transformed ratio of CoQ_6_ to PPHB_6_ abundance in WT or *fmp30*Δ strains expressing Fmp30 wild type (*FMP30 WT*), Fmp30 catalytically dead mutant (*FMP30 H265N*), NAPE-PLD wild type (*NAPE-PLD*), or mitochondrially targeted NAPE-PLD (*NAPE-PLD*^*MTS*^). **g**, Crystal structural of NAPE-PLD (gray, PDB:4QN9) with PE (red, left) and overlaid with the AlphaFold structure of Fmp30 (right). Variable alpha helix in Fmp30 is colored green. For **c, d**, and **f**, data are displayed as mean ± s.d., n=3 biologically independent samples.

Fmp30 exhibits two critical differences from NAPE-PLD: Fmp30 contains an N-terminal mitochondrial targeting sequence and a single pass transmembrane helix, both features lacking in the human protein (Fig 3a). Interestingly, only budding yeast retain both these mitochondrial localization and transmembrane domains, suggesting these proteins may have evolved alternative functions (Fig 3e). Fitting this hypothesis, neither WT nor mitochondrially-localized NAPE-PLD rescued the CoQ phenotype of *fmp30*Δ (Fig 3f). Importantly, an Fmp30 construct with a mutated metal-binding HxHxDH motif (H265N) also could not rescue this phenotype (Fig 3f), implying that Fmp30’s putative hydrolase activity is critical for its function.

The high similarity between Fmp30 and NAPE-PLD underlies the initial claim that Fmp30 is a yeast enzyme with NAPE-specific phospholipase D activity^25^, though this finding has been debated in subsequent studies due to recombinant Fmp30’s lack of *in vitro* activity^18,26^. Both proteins contain a metallo-β-lactamase domain and, expectedly given their homology, the Fmp30 AlphaFold structure aligns closely with the NAPE-PLD crystal structure (PDB:4QN9^27^) (Fig 3g). The most notable differences occur in a predicted alpha helical region that it is not resolved NAPE-PLD’s crystal structure, suggesting its flexibility (Fig 3g). In Fmp30, this flexible helix resides on top of critical residues that, in NAPE-PLD, are predicted to be essential for binding the N-acyl chain of NAPE substates (Fig 3g). These structural observations, along with differential subcellular localizations, further imply that Fmp30 and NAPE-PLD have alternative substrate specificities.

### Fmp30 possesses phospholipase D activity toward phosphatidylinositol

Since the catalytic core and active site of Fmp30 and NAPE-PLD are highly conserved, we reasoned that Fmp30 may possess phospholipase activity toward a different substrate. Surprisingly, of all phospholipids tested, only phosphatidylinositol (PI) resulted in any generation of phosphatidic acid (PA), a main product of a phospholipase D reaction (Fig 4a, b). Importantly, we observed no generation of PA with either NAPE-PLD or the catalytically-dead Fmp30 H265N mutant using PI as a substrate (Fig 4c). We reasoned that if Fmp30 performs this activity *in vivo*, then altering its expression could significantly alter the mitochondrial lipidome. We isolated highly purified mitochondria from WT, *fmp30*Δ, and a WT strain over-expressing *FMP30* driven by a high-expression promotor (GPD) (Supp Fig 4a) and performed both targeted and untargeted lipidomic analyses. Consistent with our observations at the whole cell level, deletion of *FMP30* increased the mitochondrial CoQ:PPHB ratio, while *FMP30* overexpression drove that ratio in the reverse direction (Supp Fig 4b). Deletion of *FMP30* significantly increased levels of nearly all PI species in purified mitochondria (Fig 4d); notably PI is the only phospholipid class that demonstrated such drastic changes (Supp Fig 4c). Reciprocally, overexpression of *FMP30* uniquely decreased PI species (Fig 4e, Supp Fig 4d). Both expression models exhibited the opposite effect on PA abundance, with decreased PA levels in the *FMP30* deletion and increased levels in the overexpression strain (Fig 4f).

**Figure 4.**
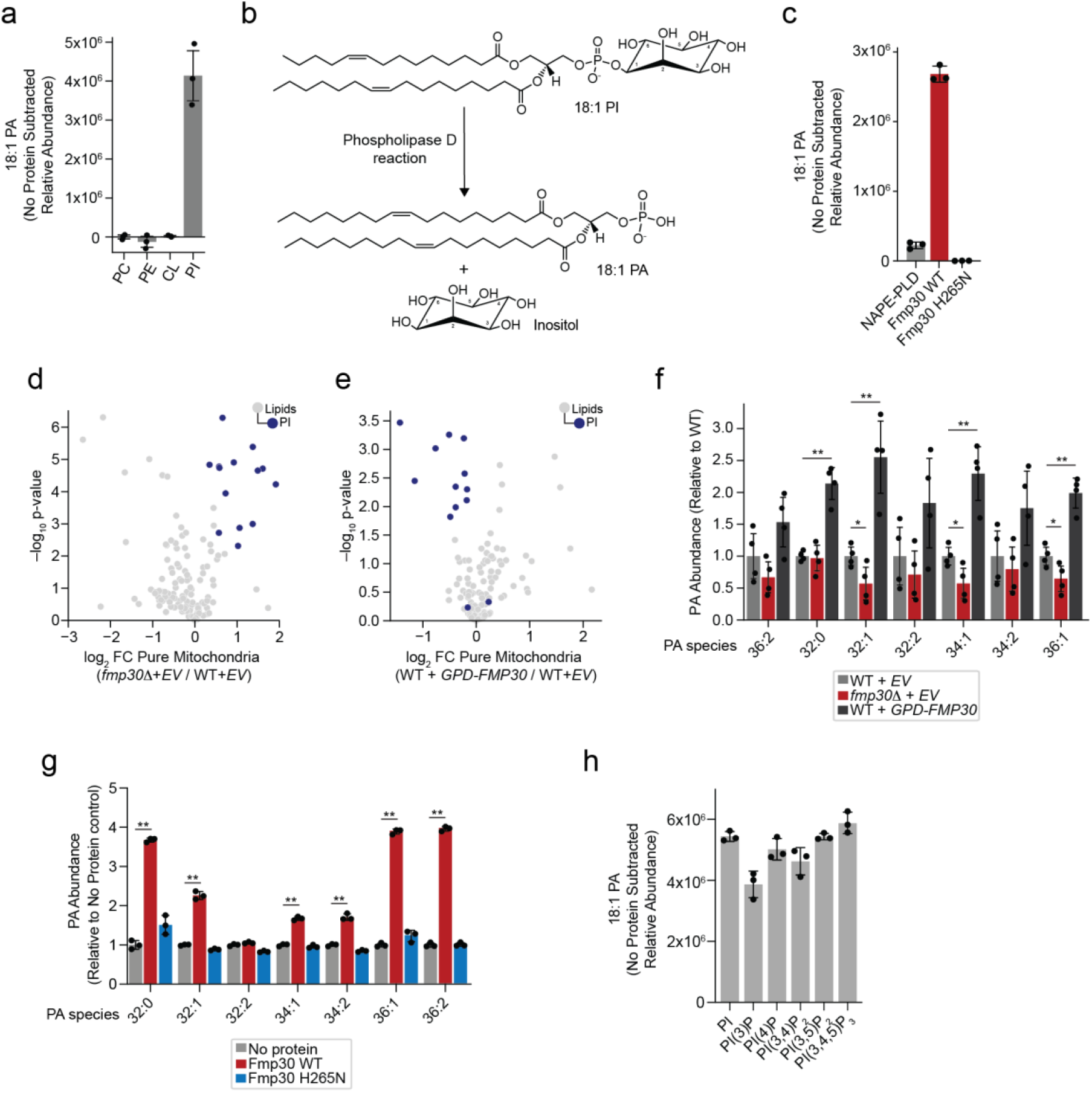
Fmp30 possesses phospholipase D activity toward phosphatidylinositol. **a**, Relative abundance of 18:1 phosphatidic acid (PA) generated after incubation of the indicated phospholipid species with recombinant Fmp30 for 1 hr. PC, phosphatidylcholine; PE, phosphatidylethanolamine; CL, cardiolipin; PI, phosphatidylinositol. **b**, Proposed phospholipase D enzymatic reaction catalyzed by Fmp30. **c**, Relative abundance of 18:1 PA after incubation of 18:1 PI with recombinant NAPE-PLD, Fmp30 WT or Fmp30 H265N for 1 hr. (**d-e)**, Relative lipid abundance in purified mitochondria from *fmp30*Δ carrying an empty vector (EV) (**d**) or WT with *FMP30* overexpression (*GPD-FMP30*) (**e**) compared to WT carrying an EV, versus statistical significance. Phosphatidylinositol (PI) species are colored in dark blue. **f**, Relative abundance of indicated phosphatidic acid (PA) species in pure mitochondria from WT carrying an EV, *fmp30*Δ carrying an EV, or WT overexpressing *FMP30* (*GPD-FMP30*). **g**, Relative abundance of indicated phosphatidic acid (PA) species following incubation of pure mitochondria from *fmp30*Δ with no protein, recombinant Fmp30 WT, or Fmp30 H265N for 1 hr. **h**, Relative abundance of 18:1 phosphatidic acid (PA) after incubation of the indicated phosphoinositol phosphate species with recombinant Fmp30 WT for 1 hr. For **a, c, g and h**, data are displayed as mean ± s.d., n=3 biologically independent samples; for **d, e** and **f**, n=4 biologically independent samples, two-sided Student’s *t*-test. \**P*<0.05, **P*<*0.01.

Collectively, these data point to PI as the native substrate of Fmp30. The activity of Fmp30 toward PI species of varying acyl-chain lengths and saturations is consistent with the substrate selectivity of NAPE-PLD; although NAPE-PLD does not hydrolyze other phospholipids, it is highly active toward NAPEs with acyl chains of various lengths^28^. Fitting with this model, incubation of recombinant Fmp30 with mitochondria isolated from *fmp30*Δ yeast significantly increased the levels of numerous PA species, with no PA observed from the Fmp30 H265N mutant (Fig 4g). Intriguingly, Fmp30 also displayed activity *in vitro* toward several phosphatidylinositol phosphate (PIP) species, suggesting that Fmp30 may have broad activity against inositol lipids (Fig 4h). How observed changes in mitochondrial PI or PIP levels in turn regulate complex Q formation or stability is an active area of investigation.

## DISCUSSION

Here, we identify the uncharacterized mitochondrial protein Fmp30 as a negative regulator of CoQ biosynthesis in *S. cerevisiae*. Using genetic manipulation, untargeted lipidomics, and biochemical approaches, we characterize Fmp30 as a phospholipase type D enzyme that hydrolyzes phosphatidylinositol and phosphoinositides to yield phosphatidic acid. We postulate that disruption of Fmp30-mediated mitochondrial phospholipid modulation results in a membrane environment that seeds complex Q formation or stabilizes CoQ domains, thus promoting CoQ biosynthesis. We have previously demonstrated that CoQ domains colocalize with ER-mitochondria contact sites^22,29^, potentially to supply biosynthetic intermediates from the ER or promote the extra-mitochondrial transport of mature CoQ. ER-mitochondria contact sites additionally have been shown to spatially coordinate and house mitochondrial fission machinery^30,31^. Furthermore, recent studies have implicated PI(4)P derived from the Golgi- and ER-mitochondria contact sites in driving mitochondrial division^32–34^. Taken together, it is plausible that deletion of *FMP30* could lead to an increase in mitochondrial PI and PIPs, which then promote ER-mitochondria contact sites and mitochondrial division. This ultimately fosters the seeding sites for CoQ domains and therefore increases biosynthetic flux.

As an alternative model, Fmp30 could be exerting its downstream effects on CoQ biosynthesis through alterations in levels of its product, phosphatidic acid. Though PA comprises only a small fraction of the total mitochondrial phospholipid pool, it plays a critical role in regulating mitochondrial morphology and dynamics^35^. PA promotes mitochondrial fusion by tethering adjacent organelles, while it also blocks mitochondrial division via inhibitory interactions with Drp1^36^. In mammals, an outer-mitochondrial membrane-localized phospholipase D, Mito-PLD, converts CL to PA^37^. Deletion of *MITO-PLD* causes mitochondrial fragmentation, presumably due to a loss of PA-mediated fusion^37^. Interestingly, a yeast strain lacking *FMP30* exhibited fragmented mitochondria, which could not be rescued by re-expression of an Fmp30 construct with mutations to its proposed catalytic residues^18^. Though Mito-PLD and Fmp30 exhibit different sub-mitochondrial localization, this phenocopying of the mitochondrial morphology defect supports a model in which Fmp30 alters mitochondrial membrane composition and CoQ domain seeding via PA production. Our current work supports both potential hypotheses: that increased CoQ biosynthesis in *fmp30*Δ could result from either increased PI/PIP levels or decreased PA levels. Work to distinguish between these mechanisms is ongoing.

Even though CoQ biology has been an active area of investigation for over 65 years, regulation of CoQ biosynthesis remains poorly characterized. Some evidence suggests CoQ biosynthesis is regulated at the transcriptional, translational, and post-translational levels, however such mechanisms are often species- and context-specific^1^. A recent study proposed the phosphatidylethanolamine methyltransferase, *CHO2*, regulates CoQ biosynthesis by controlling the mitochondrial SAM pool and thus mitochondrial methylation capacity^14^. Despite surveying the yeast deletion collection for alterations in CoQ abundance, the authors did not detect elevated CoQ levels in the strains most affected in our study (deletions in *FMP30, CHO1, UPS2, PSD1*)^14^. In our hands, we observed no increase in CoQ abundance in the *cho2*Δ strain (Fig 1b). Such inconsistencies could be attributed to differences in yeast strains or media composition. Despite these discrepancies, our work here and the work of others suggest that further investigation is required to fully explore the regulatory relationship between mitochondrial phospholipid metabolism and CoQ biosynthesis.

Despite the homology between Fmp30 and human NAPE-PLD, we present evidence that these proteins are functionally distinct. First, the two proteins have different subcellular localizations, with NAPE-PLD being membrane-associated at the Golgi, endosomes, and plasma membrane^27^, and Fmp30 localized to the mitochondrial intermembrane space, anchored to the inner mitochondrial membrane with its transmembrane domain^18^. Second, *NAPE-PLD* constructs, both its native sequence and with an engineered mitochondrial-targeting sequence, failed to rescue the CoQ phenotype of *fmp30*Δ (Fig 3f). Third, recombinant NAPE-PLD and Fmp30 displayed unique substrate specificities *in vitro*. NAPE-PLD was shown to act specifically on NAPE species, with little-to-no phospholipase D activity toward any other phospholipid class, including PI^28^. Conversely, we demonstrate that purified Fmp30 has no activity towards NAPEs (Fig 3c, d) or other phospholipid classes, only PI and PIPs (Fig 4a, h). Furthermore, the existence of NAPEs in yeast is debated^18,25^; we were unable to detect any NAPE species using high resolution LC-MS/MS in WT or *fmp30*Δ yeast at the whole cell or purified mitochondria level (data not shown). Despite high sequence similarities, enzymes within the metallo-β-lactamase super family have a diverse array of known substrates and a broad range of cellular functions^38^, therefore it is not surprising that NAPE-PLD and Fmp30 have evolved alternative substrate specificities. Our work here lays the foundation for future efforts to identify a functional mammalian counterpart in mammals.

Given the critical role of CoQ in maintaining energy homeostasis and the severe human pathologies that arise from its deficiency, it is imperative to define regulatory mechanisms that govern CoQ biosynthesis. Primary and secondary CoQ deficiencies suffer from largely ineffective therapeutic options due to the high hydrophobicity and low bioavailability of CoQ^5^. Alternative options currently being explored involve chemical bypass of dysfunctional steps in the biosynthetic process^39^ or manipulation of the pathway by small molecule probes to boost CoQ biosynthesis^40^. Identification of a human Fmp30 functional ortholog could provide an important handle for manipulating CoQ levels in a disease-relevant context. The identification and characterization of potent specific inhibitors of NAPE-PLD^41^ suggest that a mammalian ortholog of Fmp30 could represent a druggable target, thereby exposing a novel therapeutic avenue to manipulate CoQ biosynthesis *in vivo*. Collectively, our work here nominates a previously uncharacterized mitochondrial protein, Fmp30, as a negative regulator of CoQ biosynthesis. Further efforts to understand the coordination of phosphatidylinositol, phosphoinositide, and phosphatidic acid metabolism with CoQ biosynthesis, especially in a mammalian context, may enable therapeutic manipulation of this pathway.

## MATERIALS AND METHODS

### Yeast strain generation and culture conditions

The *S. cerevisiae* haploid strain BY4742 (MATa his3 leu2 lys2 ura3) was used and cultured under standard laboratory conditions. Single (*gene*Δ) and double (*gene*Δ*gene*Δ) deletion strains were generated using PCR-based homologous recombination where open reading frames were replaced with the KanMX6 or His3MX6, cassettes transformed using standard heat shock conditions and confirmed via PCR genotyping^42^. For CoQ domain imaging, the C-terminal endogenous tag on Coq9 was generated using PCR-based homologous recombination with the mNeonGreen:NatMX6 cassette.

For all whole cell lipidomic and proteomic analyses, strains from glycerol stocks were first struck out YPD plates consisting of 1% (w/v) yeast extract (‘Y’) (Research Products International), 2% (w/v) peptone (‘P’) (Research Products International), 2% (w/v) dextrose (‘D’) (Fisher) and 2% (w/v) agar (Sigma-Aldrich) and allowed to grow for 48 h at 30 °C. Starter cultures were inoculated from individual colonies in 3 mL YPD media and incubated for 24 h (30 °C, 230 rpm). Cell density was measured at OD^600^ and 1.25×10^6^ cells from each starter culture were used to inoculate 50 mL of respiratory YPG media (1% (w/v) Y, 2% (w/v) P, 0.1% D and 3% glycerol(‘G’)) in a sterile 250 mL Erlenmeyer flask. When indicated, samples were supplemented with 2 mM ethanolamine (Sigma). Samples were incubated (30 °C, 230 rpm) and 1×10^8^ cells were collected at 24 h, a timepoint that corresponds to early respiratory growth. The samples were collected by centrifugation (4,000xg, 5 min, r.t.). The supernatant was removed and the cells washed with 1 mL of sterile water. Cells were pelleted again (12,000xg, 1 min, r.t.) and the supernatant was removed. Cell pellets were snap frozen in liquid nitrogen (LN^2^) and stored at −80 °C until analysis.

### Plasmids

Yeast *fmp30*, human *NAPE-PLD*, and human *NAPE-PLD* fused with the mitochondrial targeting sequence of yeast *fmp30* were cloned into a single copy plasmid (p416) under the control of the GPD promoter. Yeast *fmp30* was cloned into an alternative p416-GPD plasmid in which the *URA3* gene was swapped for the KanMX6 cassette. For mitochondrial imaging, the Su9-BFP construct was cloned into a p416-GPD vector harboring the KanMX6 cassette. Constructs were transformed into WT and *fmp30*Δ yeast using the LiAc/SS carrier DNA/PEG method^43^. Strains harboring the p416-GPD URA3 vectors were grown in SC Ura^-^ (2% glucose, w/v) or SC Ura^-^ (0.1% glucose, 3% glycerol, w/v) as detailed above. Strains harboring the vectors containing the KanMX6 cassette were cultured as above in rich media supplemented with 200 μg/mL G418.

### Respiratory growth assays

Starter cultures (YPD, 3 mL) were inoculated with individual colonies and incubated overnight (30 °C, 230 rpm, 14–16 h). For plate reader-based assays, cells were pelleted and resuspended in respiratory media (YPG) at a density of 5×10^6^ cells/mL. Then, 100 μL of the resuspended cells were transferred to a sterile 96-well round-bottom plate (Thermo) with a Breathe-Easy cover seal (Diversified Biotech). Cultures were incubated (30 °C, 1140 rpm) in an Epoch2 plate reader (BioTek) with OD^600^ measured every 10 min.

### CoQ domain imaging

#### Fluorescence microscopy

Cells were grown as detailed above. Upon reaching mid-log phase, 1 mL of cells was collected by centrifugation at (4,000xg, 1 min) and resuspended in glucose-free media at approximately one one-hundredth of the original volume. All images were taken as z-stacks using a Zeiss LSM880 inverted laser scanning confocal microscope equipped with Zen software. Images were acquired with a 63x oil objective NA = 1.4 at room temperature.

#### Image analysis

All image analysis was performed using Fiji/ImageJ. For quantification of domain number and intensity, max projections were generated from z-stacks and used to generate masks. Briefly, max projections were background subtracted by generating a blurred image using the ‘*Gaussian blur*’ function in ImageJ with a sigma=8.00. Blurred images were subtracted from the originals using the ‘*Image calculator*’ function. Background-subtracted projections were segmented using ‘*Adjust threshold*’. Particles were auto-detected using the ‘*Analyze particles*’ function with a size range of 0.05-2.0µm^2^ and circularity range of 0.5-1.0. Particles detected in the mask were overlaid onto average intensity projections and the ‘*Measure*’ function was used to calculate integrated density. For each image, the number of cells was counted manually. For each quantification, n≥100 cells.

### Mitochondrial isolation

Starter cultures (3 mL, YPD) were inoculated with individual colonies and incubated (30 °C, 230 r.p.m., 14–16 h), then 1×10^8^ cells were diluted into 2 L of YPG in a 5L Erlenmeyer flask and incubated (30 °C, 230 r.p.m.) for 24-26 h to a final OD ≈ 3. For the overexpression studies, WT or *fmp30*Δ strains were transformed with a p416-GPD empty vector or p416-GPD-FMP30, harboring a KanMX resistance cassette in place of the *URA3* gene. In these cases, media was supplemented with 200 μg/mL G418 to maintain selection. Mitochondria were isolated as previously described^44^ . Cells were collected by centrifugation (3,000xg, 5 min, r.t.), washed with water, and centrifuged again (3,000xg, 5 min, r.t.). The wet pellet weight of the cells was determined. Cells were resuspended in 2 mL/g dithiothreitol buffer (100 mM Tris-H^2^SO^4^,10 mM dithiothreitol, pH 9.4) and shaken slowly (30 °C, 80 r.p.m., 20 min). Cells were pelleted, washed once with 7 mL/g Zymolyase buffer (1.2 M sorbitol, 20 mM potassium phosphate, pH 7.4), and resuspended in 7 mL/g Zymolyase buffer with 3 mg/g Zymolyase20T (Fisher) to generate spheroplasts. The yeast was shaken slowly (30 °C, 80 r.p.m.) for 40 min before being pelleted and washed with 7 mL/g Zymolyase buffer. Pellets were resuspended in 6 mL/g ice-cold homogenization buffer (0.6 M sorbitol, 10 mM Tris-HCl, 1 mM phenylmethylsulfonyl fluoride (PMSF), 0.2% (wt/vol) fatty-acid-free bovine serum albumin (BSA), pH 7.4). Spheroplasts were homogenized using 20 strokes of a tight-fitting glass-Teflon homogenizer and diluted two-fold with homogenization buffer. The homogenate was centrifuged (1,500xg, 5 min, 4 °C) to pellet cell debris and nuclei. The supernatant was centrifuged (4,000xg, 5 min, 4 °C) to pellet additional debris. Crude mitochondria were isolated by centrifuging the supernatant (12,000xg, 15 min, 4 °C) and resuspended in SEM (250 mM sucrose, 1 mM EDTA, 10 mM MOPS-KOH, pH 7.2). For isolation of pure mitochondria, crude mitochondrial fractions in SEM were incubated with 10 μg trypsin (sequencing grade, Promega) and rotated end-over-end overnight (16 h, 4 °C) to disrupt proteinaceous organelle contact tethers^45^. On the following day, digested samples were pelleted by centrifugation (15,000xg, 7 min, 4 °C). Pelleted material was resuspended in 900 μL SEM buffer containing 1 mM PMSF to deactivate trypsin. Resuspended material was pelleted (15,000xg, 7 min, 4 °C) and this was repeated once more. Pelleted crude mitochondria were resuspended in 200 μL SEM+PMSF and then added to a freshly prepared sucrose gradient (bottom to top: 1.5mL 60% sucrose, 4 mL 32% sucrose, 1.6 mL 23% sucrose, and 1.4 mL 15% sucrose) for separation by ultracentrifugation (134,000xg, 1 h, 4 °C). Enriched mitochondrial samples were recovered at the 32−60% interface and diluted with 30 mL SEM. Mitochondria were pelleted (15,000xg, 10 min, 4 °C) and resuspended in fresh SEM. Mitochondrial protein content was quantified by a BCA protein assay (Thermo). Pure mitochondrial samples were snap frozen as 75 μg aliquots for lipidomic analysis.

### Cross-linking mass spectrometry

Crude mitochondria were isolated from yeast as described above and subjected to chemical cross-linking (1 mg mitochondria, 0.5 mM DSSO (Thermo, catalogue no. A33433), 1 hour, r.t.). Cross-linking was quenched with 100 mM Tris pH 8.0 followed by centrifugation (15,000xg, 5 min, 4 °C). The mitochondria were then solubilized with 50 mM imidazole, 500 mM 6-hexaminocaproic acid, 1 mM EDTA and 1 g/g digitonin. The bait protein and cross-linked interactors were then enriched by mNeonGreen immunoprecipitation (IP) using magnetic mNeonGreen-Trap agarose beads (Chromotek, catalogue no. nta-20), washed and subjected to on-bead tryptic digest. The on-bead cross-linked proteins were denatured with 2 M urea in 200 mM Tris pH 8.0, then reduced with 5 mM DTT for 30 min at 56 °C and alkylated with 15 mM iodoacetamide for 30 min at r.t. in the dark. The proteins on-bead were digested overnight at 37 °C with 1 μg trypsin (Promega, catalogue no. V5113). The digested supernatant was acidified with 10% TFA to a pH of 2 and desalted with 10 mg StrataX solid phase extraction columns (Phenomenex), then dried under vacuum using a SpeedVac (Thermo Scientific) and stored at −80 °C until MS analysis.

Samples were resuspended in 0.2% formic acid and subjected to LC–MS analysis. LC separation was performed using the Thermo Ultimate 3000 RSLCnano system. A 15 cm EASY-Spray PepMap RSLC C18 column (150 mm × 75 μm, 3 μm) was used at 300 nL/min flow rate with a 90 min gradient using mobile phase A consisting of 0.1% formic acid in H^2^O, and mobile phase B consisting of 0.1% formic acid in ACN/H^2^O (80/20, v/v). An EASY-Spray source was used and the temperature was 35 °C. Each sample run was held at 4.0% B for 5 min and increased to 50% B over 65 min, followed by 8 min at 95% B and back to 4% B for equilibration for 10 min. An Acclaim PepMap C18 HPLC trap column (20 mm × 75 μm, 3 μm) was used for sample loading. MS detection was performed with a Thermo Exploris 240 Orbitrap mass spectrometer in positive mode. The source voltage was set to 1.8 kV, ion transfer tube temperature was set to 275 °C, RF lens was at 70%. Full MS spectra were acquired from m/z 350 to 1,400 at the Orbitrap resolution of 60,000, with the normalized automatic gain control target of 300% (3 × 10^6^). Data-dependent acquisition (DDA) was performed for the top 20 precursor ions with the charge state of 2–6 and an isolated width of 2. Intensity threshold was 5 × 10^3^. Dynamic exclusion was 30 s with the exclusion of isotopes. Other settings for DDA include Orbitrap resolution of 15,000 and high-energy collision-induced dissociation energy of 30%.

Raw files were analyzed by SequestHT search engine incorporated in Proteome Discoverer v.2.5.0.400 software against yeast databases downloaded from Uniprot. Label-free quantification was enabled in the searches. The resulting data were analyzed by Perseus v.1.6.15.0 software^46^ .

### LC-MS/MS proteomics

#### Yeast growth

Yeast cultures were grown as described previously for respiration, then 1×10^8^ cells were collected, snap-frozen in LN^2^, and stored at −80 °C.

#### Lysis and digestion

Yeast pellets were removed from −80 °C and resuspended in lysis buffer (6 M guanidine hydrochloride,100 mM Tris). The samples were then boiled at 100 °C for 5 min and subjected to probe sonication (2 rounds of 10 s on, 20 s off, 25% amplitude). Methanol was added to each sample to 90% concentration, and the samples were centrifuged at 9,000xg for 30 min to precipitate proteins. After precipitation, the supernatant was discarded from each sample, and the protein pellets were allowed to air-dry. The dried pellets were resuspended in digestion solution (8 M urea, 10 mM TCEP, 40 mM CAA, 100 mM Tris pH 8.0), and the samples were sonicated in the bath sonicator to facilitate re-solubilization. Trypsin (Promega, catalogue no. V5113) was added to each sample in a 50:1 protein/enzyme ratio before they were incubated at 37 °C overnight. The peptides were dried under vacuum using a SpeedVac and resuspended in 0.1% TFA. High pH fractionation was performed according to the manufacturer’s protocol (Pierce High pH Reversed-Phase Peptide Fractionation Kit, Thermo Scientific). Eight fractions were collected and dried under vacuum (Thermo Scientific).

#### LC-MS/MS proteomics data acquisition

Peptides were resuspended in 0.2% formic acid, and the concentration of each sample was determined using a NanoDrop One spectrophotometer (Thermo Scientific). LC separation was performed using the Thermo Ultimate3000 RSLC nano system. A 15 cm EASY-Spray PepMap RSLC C18 column (Thermo, 150 mm × 75 μm, 3 μm) was used at 300 nL/min flow rate with an Acclaim PepMap C18 HPLC trap column (Thermo, 20 mm × 75 μm, 3 μm) for sample loading. For each sample run, the temperature was held at 35 °C for a 120 min gradient that consisted of 4% B for 5 min and increased to 30% B over 100 min, followed by 5 min at 99% B and back to 4% B for equilibration for 10 min. Mobile phase A consisted of 0.1% FA in water, and mobile phase B consisted of 0.1% FA in 80% (v/v) ACN and 20% (v/v) water. MS detection was performed with a Thermo Exploris 240 Orbitrap mass spectrometer with an EASY-Spray source operating in positive mode. The source voltage was 1.8 kV, the ion transfer tube temperature was set to 275 °C and the RF lens at 70%. Full MS spectra were acquired from m/z 350 to 1,400 at the Orbitrap resolution of 60,000, with a normalized AGC target of 300% (3 × 10^6^). Data-dependent acquisition was performed with a 3 s duty cycle with a charge state of 2–6, an isolation window width of 2 and an intensity threshold of 5 × 10^3^. Dynamic exclusion was 20 s with the exclusion of isotopes. Other settings for data-dependent acquisition were an Orbitrap resolution of 15,000 and higher energy collisional dissociation energy of 30%.

#### Data analysis

Raw files were analyzed by the SequestHT Search Engine incorporated in Proteome Discoverer v.2.5.0.400 software against yeast databases downloaded from Uniprot. Label-free quantification was enabled in the searches.

### LC-MS/MS Lipidomics

#### Yeast growth for *de novo* CoQ measurements

Starter cultures (SC 2% glucose, w/v, 3 mL) were inoculated with individual colonies and incubated overnight (30 °C, 230 rpm, 14–16 h). Next, 50 mL cultures in SC pABA^−^ (2% glucose, w/v) were inoculated with 2.5×10^7^ cells and incubated (30 °C, 230 rpm) until reaching an OD^600^ ∼ 2. The cells were then centrifuged (3,000xg, 3 min, r.t.) and resuspended in SC pABA^−^ media (3% glycerol, w/v) containing ^13^C_6_-4HB (50 μM, Sigma) and incubated (30 °C, 230 rpm). At 2, 4 and 6 h after the media swap, 1×10^8^ cells were collected, snap-frozen in LN^2^, and stored at −80 °C.

#### Lipid Extraction

Lipids from cell pellets or pure mitochondrial samples were extracted using MTBE (Sigma-Aldrich). Frozen cell pellets were resuspended in 225 μL 100% LC-grade methanol (Fisher), containing 1 μM CoQ^8^ (Avanti Lipids) as an internal standard. Glass beads (100 μL, 0.5 mm; BioSpec) were then added and the samples were vortexed using a Vortex Genie for 10 min (3,000 rpm, 4 °C) to lyse the cells. Next, 187.5 μL of water and 750 μL of MTBE were added to each sample and the tubes were vortexed again for 3 min (3,000 rpm, 4 °C). To separate the layers, samples were centrifuged for 3 min (1,000xg, 4 °C). The organic (top) layer was removed into a separate microcentrifuge tube and a new 750 μL of MTBE was added. Organic extraction was repeated a second time with the second MTBE layer added to the first. Samples were dried by vacuum centrifugation and resuspended in 50 μL of 20 mM ammonium acetate in 78% (v/v) methanol, 20% IPA (Sigma-Aldrich) and 2% water.

#### Untargeted LC-MS/MS lipidomics data acquisition

A Vanquish Horizon UHPLC system (Thermo Scientific) connected to an Exploris 240 Orbitrap mass spectrometer (Thermo Scientific) was used for LC-MS analysis. A Waters Acquity CSH C18 column (100 mm × 2.1 mm, 1.7 μm) was held at 50 °C with the flow rate of 0.4 mL/min for lipid separation. A Vanquish binary pump system was employed to deliver mobile phase A, consisting of 5 mM ammonium acetate in ACN/H^2^O (70/30, vol/vol) containing 250 μL/L acetic acid, and mobile phase B consisting of 5 mM ammonium acetate in IPA/ACN (90/10, vol/vol) containing 250 μL/L acetic acid. The gradient was set as follows: B was at 2% for 2 min and increased to 30% over the next 1 min, then further ramped up to 50% within 1 min and to 85% over the next 14 min, and then raised to 99% over 1 min and held for 7 min, before re-equilibration for 4 min at 2% B. Five microliters of the sample were injected by a Vanquish Split Sampler HT autosampler (Thermo Fisher Scientific), while the autosampler temperature was kept at 4 °C. Two microliters of the sample were injected by a Vanquish Split Sampler HT autosampler (Thermo Fisher Scientific), while the autosampler temperature was kept at 4 °C. Samples were ionized by a heated ESI source with a vaporizer temperature of 200 °C. The sheath gas was set to 25 units, auxiliary gas to 15 units and the sweep gas to 2 unit. For untargeted discovery lipidomics, the MS was operated in polarity switching mode with the spray voltage set to 3,500 V for positive mode and 2,500 V for negative mode. The inlet ion transfer tube temperature was kept at 300 °C with 70% RF lens. Full MS1 scans were acquired at 22,500 resolution (at 200 m/z), a max ion accumulation time of 100 ms and with a scan range of m/z 200– 1,600. MS2 scans (top 3) were acquired at 30,000 resolution (at 200 m/z), max ion accumulation time of 50 ms, a 1.0 m/z isolation window, stepped NCE at 20%, 30% and 40%, and a 10.0 s dynamic exclusion. Automatic gain control (AGC) targets were set to standard mode for both MS1 and MS2 acquisitions.

#### Targeted LC-MS/MS lipidomics data acquisition

A Vanquish Horizon UHPLC system (Thermo Scientific) connected to an Exploris 240 Orbitrap mass spectrometer (Thermo Scientific) was used for LC-MS analysis. A Waters Acquity CSH C18 column (100 mm × 2.1 mm, 1.7 μm) was held at 35 °C with the flow rate of 0.3 mL/min for lipid separation. A Vanquish binary pump system was employed to deliver mobile phase A, consisting of 5 mM ammonium acetate in ACN/H^2^O (70/30, vol/vol) containing 250 μL/L acetic acid, and mobile phase B consisting of 5 mM ammonium acetate in IPA/ACN (90/10, vol/vol) containing 250 μL/L acetic acid. The gradient was set as follows: B was at 2% for 2 min and increased to 30% over the next 3 min, then further ramped up to 50% within 1 min and to 85% over the next 14 min, and then raised to 99% over 1 min and held for 4 min, before re-equilibration for 5 min at 2% B. Five microliters of the sample were injected by a Vanquish Split Sampler HT autosampler (Thermo Fisher Scientific), while the autosampler temperature was kept at 4 °C. Samples were ionized by a heated ESI source with a vaporizer temperature of 200 °C. The sheath gas was set to 40 units, auxiliary gas to 8 units and the sweep gas to 1 unit. The ion transfer tube temperature was kept at 300 °C with a 70% RF lens. The spray voltage was set to 3,500 V for positive mode and 2,500 V for negative mode. Targeted acquisition was performed using parallel reaction monitoring mode with polarity switching, targeting scans to CoQ^6^ (m/z 591.4408), ^13^C^6^-CoQ^6^ (m/z 597.4609), and CoQ^8^ (m/z 727.5660) in positive polarity, and PPHB^6^ (m/z 545.4000) and ^13^C^6^-PPHB^6^ (m/z 551.4201) in negative polarity. MS acquisition parameters include resolution of 15,000, an isolation window of 2 m/z, stepped HCD energies of 20%, 40% and 60% for positive mode or 25%, 30%, and 40% for negative mode, standard AGC target and auto maximum ion injection time.

#### Data analysis

For untargeted lipidomic analyses, LC–MS files were processed using Compound Discoverer 3.2 (Thermo Scientific) and LipiDex^47^. All peaks with a 1.4–23 min retention time and 100–5,000 Da MS1 precursor mass were aggregated into compound groups using a 10 ppm mass tolerance and 0.4 min retention time tolerance. Peaks were excluded if peak intensity was less than 2×10^6^, peak width was greater than 0.75 min, signal-to-noise ratio was less than 1.5 or intensity was <3-fold greater than the blank. MS2 spectra were searched against an in silico generated spectral library^48^. Spectra matches with a dot product score >500 and reverse dot product score >700 were retained for further analysis. Lipid MS/MS spectra that contained <75% interference from coeluting isobaric lipids, eluted within a 3.5 median absolute retention time deviation of each other and were found within at least four processed files were used for identification at the individual fatty acid substituent levels of structural resolution. If individual fatty acid substituents were unresolved, then identifications were made with the sum of the fatty acid substituents. Lipid identifications were filtered with the Degreaser module within LipidDex2 (v0.1.0)^49^, based on retention time modelling. The retention time tolerance used was 0.5 min. Unreliable identifications were discarded.

Targeted quantitative analysis of all acquired compounds was processed using TraceFinder 5.1 (Thermo Scientific) with a mass accuracy of 5 ppm. The result of peak integration was manually examined.

### Immunoblotting

#### Antibodies

Primary antibodies used in this study include anti-Kar2 (SCBT sc-33630, 1:5000), and anti-Cit1^50^ (custom made at Biomatik, 1:4000). Secondary antibodies include anti-mouse immunoglobulin-G (IgG) horseradish peroxidase (HRP)-linked (Cell Signaling Technology #7076, 1:5,000) and anti-rabbit IgG HRP-linked (Cell Signaling Technology #7074, 1:5,000).

#### Mitochondrial isolation western blotting

Spheroplast, crude mitochondria, and pure mitochondria samples were collected throughout the mitochondrial isolation process. Samples were solubilized in RIPA buffer and protein concntrations were determined by BCA protein assay (Thermo). Protein (10 μg) was loaded onto NuPAGE 4–12% Bis-Tris gels (Thermo) and separated (200 V, 35 min). Proteins were transferred to a polyvinylidene fluoride (PVDF) membrane (Sigma) and blocked with 5% non-fat dry milk (NFDM) in tris-buffered saline with 0.1% Tween-20 (TBST) (1 h, r.t.). The membranes were then probed with primary antibodies diluted in 5% NFDM in TBST (overnight, 4 °C). Membranes were washed three times with TBST and then probed with secondary antibodies diluted in 5% NFDM in TBST (1 h, r.t.). Membranes were washed three times with TBST and developed by enhanced chemiluminescence (ECL) using the SuperSignal West Dura substrate (Thermo, 34075). Developed membranes were imaged on a ChemiDoc system running Image Lab Touch Software 3.0.1.14 (Bio-Rad).

### Protein purification

#### NAPE-PLD

NAPE-PLD was purified as described previously^27^ with some modifications. *E. coli* BL21-CodonPlus (DE3)-RIPL cells (Thermo) harboring plasmid with His^8^-MBP-NAPE-PLD NΔ46 were grown in a 2L LB culture at 37 °C to an OD^600^ of 0.6 and expression was induced with 1 mM isopropyl-B-D-thiogalactoside (IPTG) at 22 °C for 16 h. The cells were harvested by centrifugation (5000xg, 10 min, 4 °C) and stored at -20 °C. The cell pellet was resuspended in 35 mL Lysis Buffer (20 mM HEPES pH 7.4, 200 mM NaCl, 0.1% Triton X-100, 1 Roche protease COmplete inhibitor tablet, 1 mg/mL lysozyme) and lysed via sonication on ice (5 rounds of 20 s on, 60 s off, 75% amplitude). The lysate was clarified by centrifugation (15,000xg, 30 min, 4 °C). The cleared lysate was mixed with amylose resin (New England Biolabs) and incubated on a rotor (2 h, 4 °C). The resin was washed five times with Wash Buffer (20 mM HEPES pH 7.4, 200 mM NaCl, 1mM EDTA, 0.05% Triton X-100) and the protein was eluted with Wash Buffer containing 10 mM maltose. The protein concentration was determined by BCA protein assay (Thermo) and protein was snap frozen and stored at -80 °C.

#### Fmp30

E. coli OverExpress C41(DE3) cells (Sigma) harboring plasmid with His^8^-MBP-Fmp30 NΔ76 (wild type or H265N) were grown in a 2L LB culture at 37 °C to an OD^600^ of 0.6 and expression was induced with 1 mM IPTG at 22 °C for 16 h. The cells were harvested by centrifugation (5000xg, 10 min, 4 °C) and stored at -20 °C. The cell pellet was resuspended in 35 mL Lysis Buffer (50 mM HEPES pH 7.4, 250 mM NaCl, 5% glycerol, 0.5% DDM, 1 Roche protease COmplete inhibitor tablet, 1 mg/mL lysozyme) and lysed via sonication on ice (5 rounds of 20 s on, 60 s off, 75% amplitude). The lysate was clarified by centrifugation (15,000xg, 30 min, 4 °C). The cleared lysate was mixed with amylose resin (New England Biolabs) and incubated on a rotor (2 h, 4 °C). The resin was washed five times with Wash Buffer (50 mM HEPES pH 7.4, 250 mM NaCl, 5% glycerol, 0.05% DDM) and the protein was eluted with Wash Buffer containing 10 mM maltose. The eluted protein was concentrated with a MW-cutoff spin filter (50 kDa MWCO) and subjected to size exclusion chromatography on a HighLoad 16/600 Superdex 200 pg (GE Healthcare). Fractions containing MBP-Fmp30, assessed by SDS-PAGE, were pooled and concentrated with a MW-cutoff spin filter. The protein concentration was determined by BCA protein assay (Thermo) and protein was snap frozen and stored at - 80 °C.

### Enzyme activity assays

#### Bis-pNPP phosphodiesterase assay

Phosphodiesterase activity was measured via hydrolysis of the phosphodiester bond of the generic colorimetric reagent bis-pNPP. Protein (10 μg) was incubated with bis-pNPP (25 mM) in 100 μL assay buffer (50 mM Tris pH 8.5, 0.1% DDM, 1 mM MnCl^2^, 1 mM CoCl^2^). The reaction was monitored by increase in absorbance in Epoch2 plate reader (BioTek), with OD^410^ measured every minute.

#### Enzyme assay and lipid extraction

All lipid substrates (18:1 PE-N-18:1, 18:1 PA, 18:1 (Δ9-Cis) PE, 18:1 (Δ 9-Cis PC), 18:1 CL, 18:1 PI, 18:1 PI(3)P, 18:1 PI(4)P, 18:1 PI(3,4)P^2^, 18:1 PI(3,5)P^2^, 18:1 PI(3,4,5)P^3^) were purchased from Avanti. The activity of recombinant MBP-Fmp30 NΔ76 or MBP-NAPE-PLD NΔ46 was measured by incubating the protein (10 μg) with 1 μM of the substrate in 187.5 μL assay buffer (50 mM Tris pH 8.5, 0.1% DDM, 1 mM MnCl^2^, 1 mM CoCl^2^) for 1 h at 37 °C. For reactions using mitochondria purified from *fmp30*Δ as a substrate, mitochondria (25 μg) were first solubilized by incubation with 1 g/g DDM (detergent/protein w/w) in solubilization buffer (50 mM NaCl, 50 mM imidazole, 2 mM 6-aminohexanoic acid, 1 mM EDTA, pH 7.0). Solubilized mitochondria were diluted in assay buffer with 20 μg protein and incubated for 1 hr at 37 °C. The reactions were terminated by adding 225 μL of methanol containing 100 pmol [^2^H^4^]-palmitoylethanolamide (Caymen Chemical) as the internal standard. Lipid extraction was conducted with MTBE as described above.

#### Targeted LC-MS/MS lipidomics data acquisition

A Vanquish Horizon UHPLC system (Thermo Scientific) connected to an Exploris 240 Orbitrap mass spectrometer (Thermo Scientific) was used for LC-MS analysis. A Waters Acquity CSH C18 column (100 mm × 2.1 mm, 1.7 μm) was held at 55 °C with the flow rate of 0.3 mL/min for lipid separation. A Vanquish binary pump system was employed to deliver mobile phase A, consisting of 2.5 mM ammonium bicarbonate in IPA/ACN/H^2^O (15/35/50, vol/vol), and mobile phase B consisting of 2.5 mM ammonium bicarbonate in IPA/ACN/H^2^O (70/25/5, vol/vol). The gradient was set as follows: B started at 20% and increased to 45% over the next 2.5 min, then further ramped up to 65% within 7 min and to 85% over the next 3 min, and then raised to 99% over 6 min and held for 3 min, before re-equilibration for 4 min at 20% B. Five microliters of the sample were injected by a Vanquish Split Sampler HT autosampler (Thermo Fisher Scientific), while the autosampler temperature was kept at 4 °C. Samples were ionized by a heated ESI source with a vaporizer temperature of 200 °C. The sheath gas was set to 25 units, auxiliary gas to 15 units and the sweep gas to 5 unit. The ion transfer tube temperature was kept at 300 °C with a 70% RF lens. The spray voltage was set to 2,600 V for positive mode and 2,100 V for negative mode. Targeted acquisition was performed using parallel reaction monitoring mode with polarity switching, targeting scans to 32:0 PA (m/z 647.4657), 32:1 PA (m/z 645.4501), 32:2 PA (m/z 643.4344), 34:1 PA (m/z 673.4814), 34:2 PA (m/z 671.4657), 36:1 PA (m/z 701.5127) in negative polarity and [^2^H^4^]-palmitoylethanolamide (m/z 304.3148), 18:1 PE-N-18:1 (m/z 1008.7991) and 18:1 NAE (m/z 326.3054) in positive polarity. MS acquisition parameters include resolution of 30,000, an isolation window of 2 m/z, stepped HCD energies of 25%, 30%, and 40%, standard AGC target and auto maximum ion injection time.

#### Data analysis

Targeted quantitative analysis of all acquired compounds was processed using TraceFinder 5.1 (Thermo Scientific) with a mass accuracy of 5 ppm. The result of peak integration was manually examined.

## ACKNOWLEDGEMENTS

We would like to thank the Pagliarini Lab for their helpful feedback and discussion throughout the duration of this study. This work was supported by NIH award R35 GM131795 (D.J.P.), as well as funds from the BJC Investigator Program (to D.J.P.). D.J.P. is an investigator of the Howard Hughes Medical Institute.

## AUTHOR CONTRIBUTIONS

Z.N.B., R.M.G., and D.J.P. led the conception, design, and execution of this study and wrote this manuscript. S.W.R. performed the confocal imaging experiments. Z.N.B. and R.M.G. performed all other experiments and analyses.

## SUPPLEMENTAL FIGURES

**Supplemental Figure 1.**
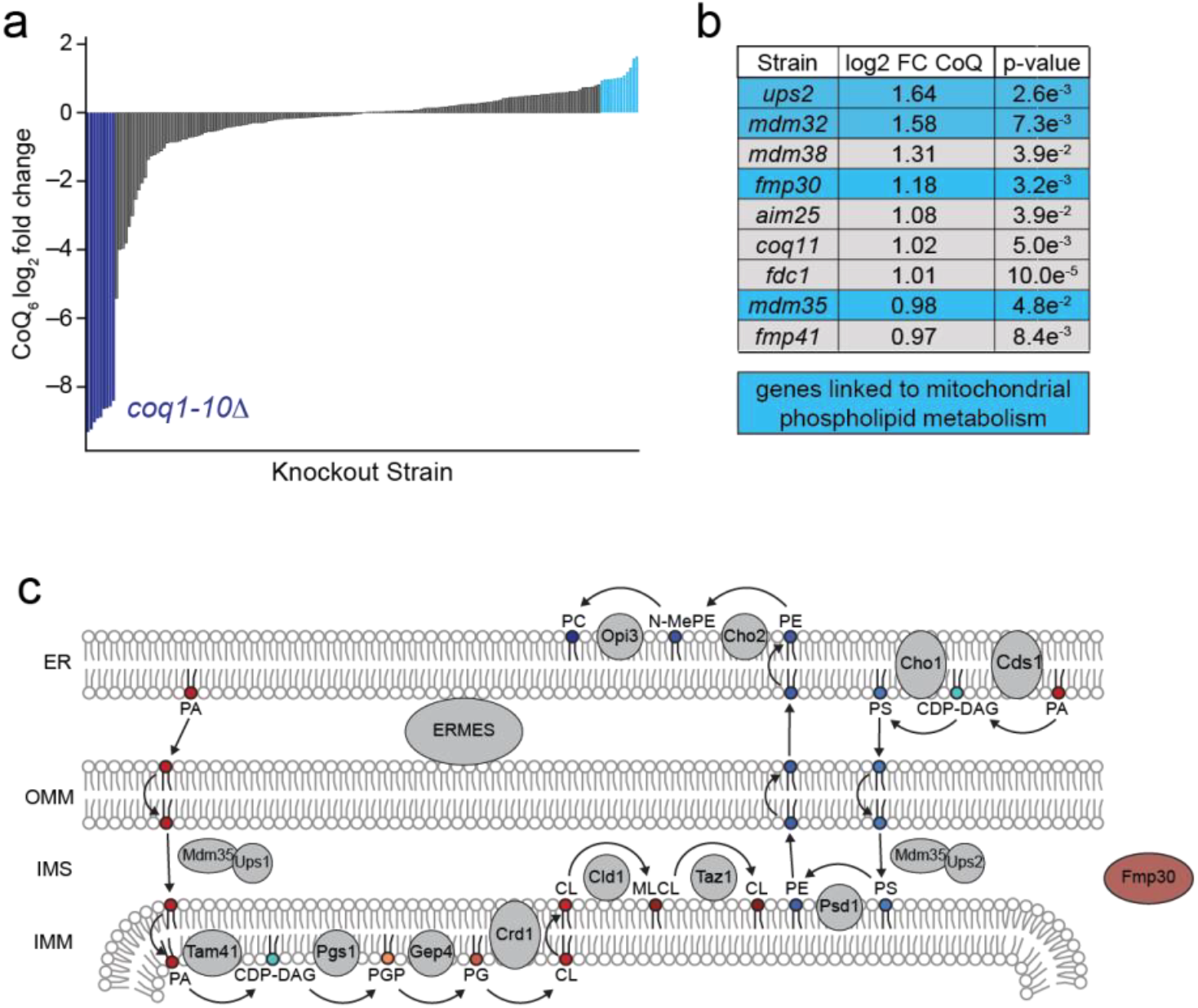
The Y3K dataset nominates potential regulators of CoQ biosynthesis. **a**, Rank ordering of gene deletion strains in the Y3K dataset^15^ based on their CoQ_6_ abundance. Strains with deletion in genes involved in CoQ biosynthesis are highlighted in dark blue. Strains with increased CoQ abundance are highlighted in light blue. **b**, Deletion strains from Y3K ranked by highest increase in CoQ abundance compared to WT. Log_2_ fold change derived from mean abundances of n=3 biologically independent samples, *p*-value from two-sided Student’s *t*-test. Genes linked to mitochondrial phospholipid biosynthesis are highlighted in light blue. **c**, Schematic of mitochondrial phospholipid biosynthetic pathways. PA, phosphatidic acid; CDP-DAG, cytidine diphosphate diacylglycerol; PGP, phosphatidylglycerol phosphate; PG, phosphatidylglycerol; CL, cardiolipin; MCLC, monolysocardiolipin; PE, phosphatidylethanolamine; PS, phosphatidylserine; N-Me-PE, N-methyl-phosphatidylethanolamine; PC, phosphatidylcholine.

**Supplemental Figure 2.**
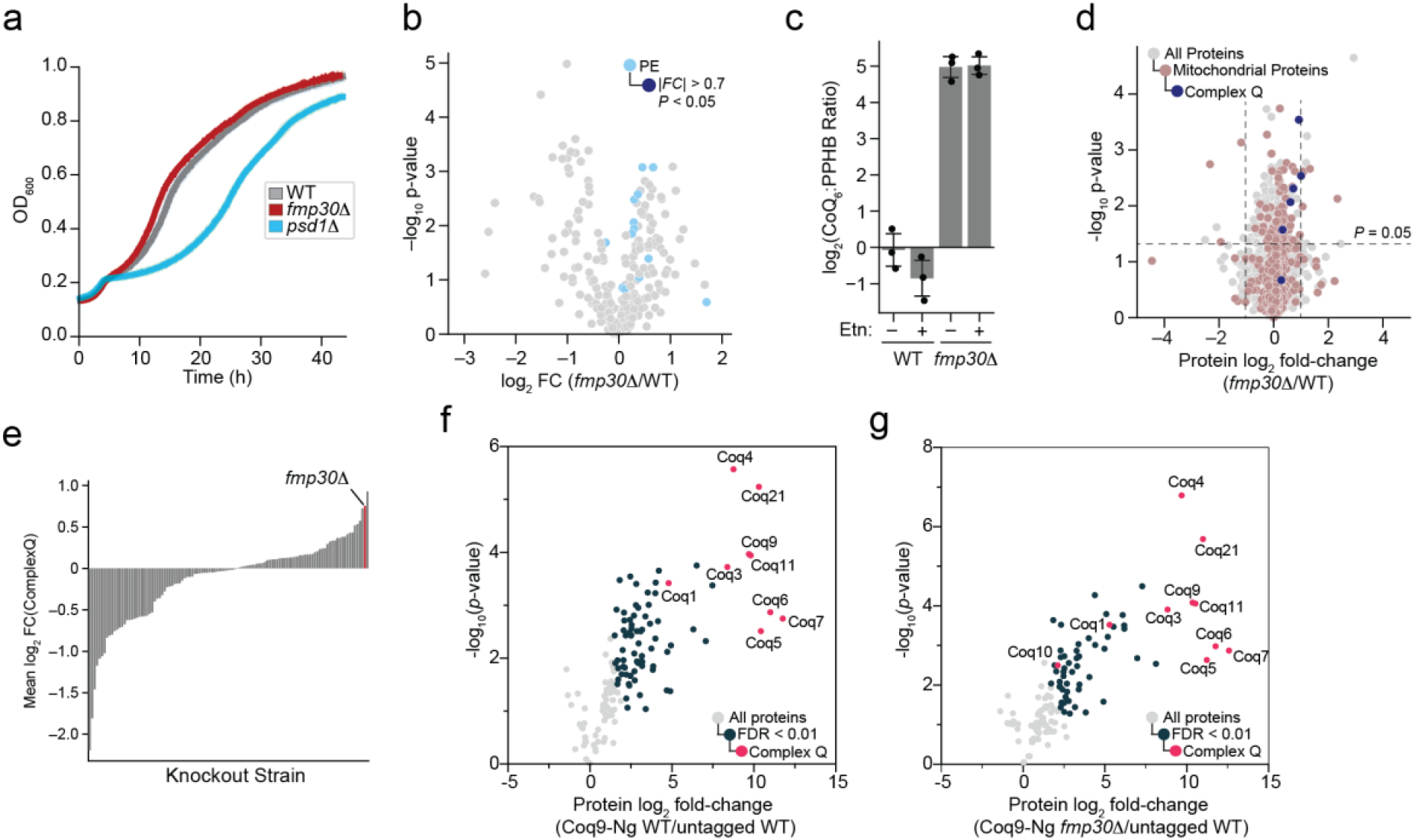
*FMP30* deletion increases the abundance of complex Q while not changing the complex Q composition. **a**, Respiratory growth assay of WT, *fmp30*Δ, and *psd1*Δ yeast in YPG media. **b**, Relative lipid abundance versus statistical significance of *fmp30*Δ compared to WT. Quantified phosphatidylethanolamine (PE) species are colored light blue, with PE species that are significantly changed (|FC| >0.7, P < 0.05; two-sided Student’s t-test) highlighted dark blue. All other lipid species are colored grey. **c**, Log_2_ transformed ratio of CoQ_6_ to PPHB_6_ abundance in WT and *fmp30*Δ following supplementation with 2 mM ethanolamine or vehicle. **d**, Relative protein abundances of *fmp30*Δ compared to WT versus statistical significance. Mitochondrial proteins are highlighted in light red, complex Q proteins are highlighted in dark blue. **e**, Rank ordering of gene deletion strains in the Y3K dataset^15^ based on their mean complex Q protein abundance. (**f**-**g**), Relative protein abundance versus statistical significance of proteins enriched following mitochondrial crosslinking and mNeonGreen immunoprecipitation of endogenously expressed Coq9-mNeonGreen in a WT strain (**f**) or *fmp30*Δ strain (**g**) compared to a WT strain expressing no tagged bait. n = 3 biological replicates, two-sided Student’s *t*-test.

**Supplemental Figure 3.**
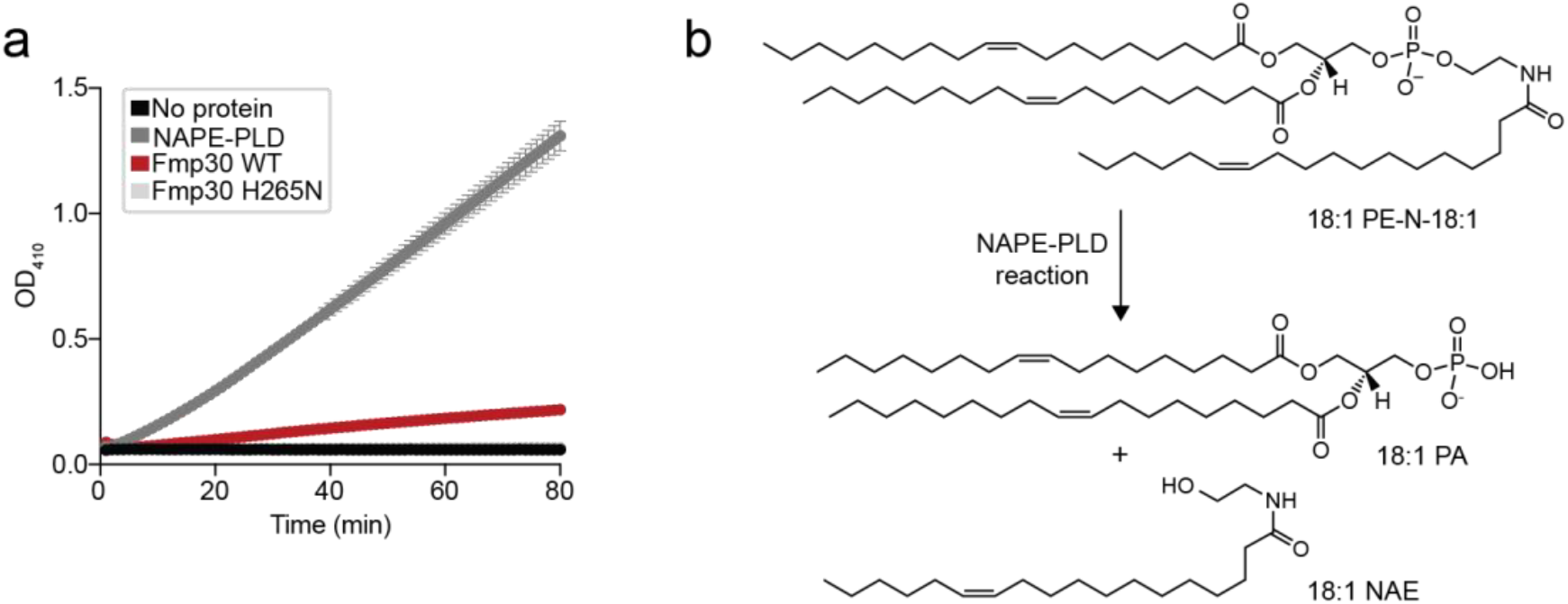
Biochemical validation of NAPE-PLD and Fmp30 phosphodiesterase activity. **a**, Phosphodiesterase activity of NAPE-PLD, Fmp30 WT, and Fmp30 H265N as measured by colorimetric reaction with the generic substrate bis-pNPP. **b**, Phospholipase D enzymatic reaction catalyzed by NAPE-PLD. PE-N, n-acyl phosphatidylethanolamine; PA, phosphatidic acid; NAE, N-acylethanolamide.

**Supplemental Figure 4.**
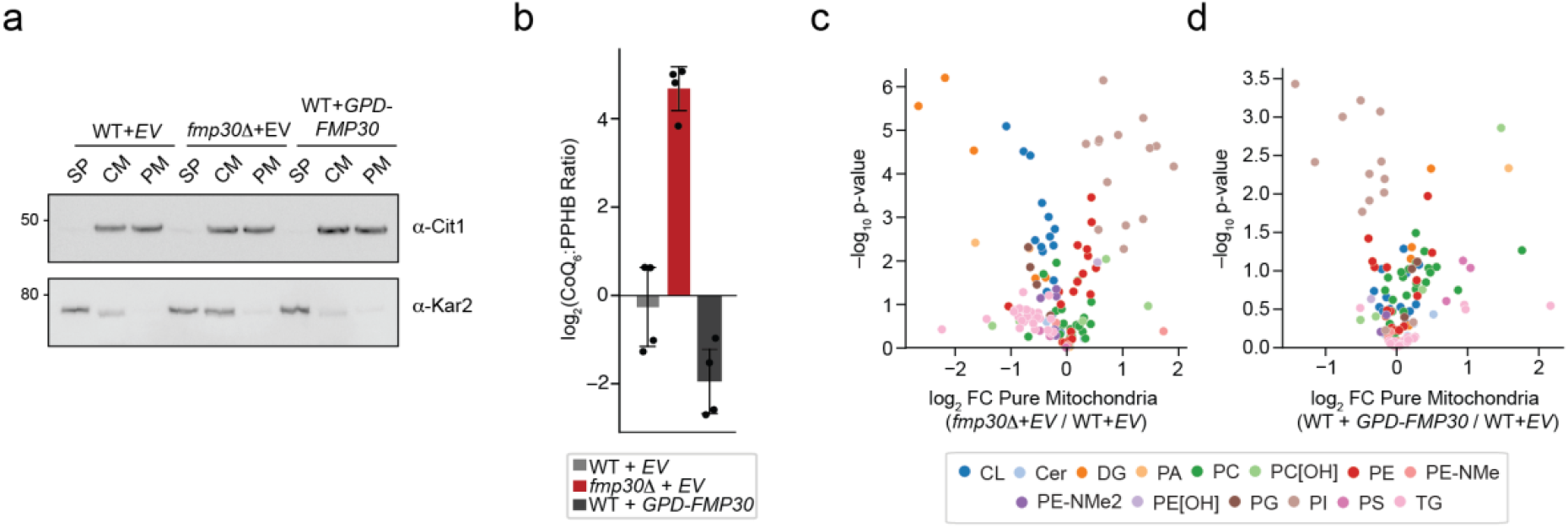
Validation and characterization of pure mitochondria with *FMP30* deletion and overexpression. **a**, Western blot of subcellular samples derived from fractionated WT + *EV, fmp30*Δ + *EV*, and *fmp30*Δ + *GPD-FMP30* yeast. SP, spheroplast; CM, crude mitochondria; PM, pure mitochondria. Kar2, endoplasmic reticulum; Cit1, mitochondria. A representative Western blot from 4 independent experiments. **b**, Log_2_ transformed ratio of CoQ_6_ to PPHB_6_ abundance in mitochondria isolated from WT + *EV, fmp30*Δ + *EV*, and *fmp30*Δ + *GPD-FMP30* yeast. Data normalized to WT + *EV*. (**c**-**d**), Data from **Fig 4 d-e** showing relative lipid abundance in purified mitochondria from *fmp30*Δ + *EV* (**c**) or WT + GPD-*FMP30* (**d**) compared to WT + *EV*, versus statistical significance. Points are color-coded according to lipid class. For **b**-**d**, n = 4 independent biological experiments.

## REFERENCES

1. Guerra, R. M. & Pagliarini, D. J. Coenzyme Q biochemistry and biosynthesis. Trends Biochem Sci 5, 463–476 (2023).

2. Wang, Y., Lilienfeldt, N. & Hekimi, S. Understanding coenzyme Q. Physiol. Rev. 104, 1533–1610 (2024).

3. Alcázar-Fabra, M., Trevisson, E. & Brea-Calvo, G. Clinical syndromes associated with Coenzyme Q10 deficiency. Essays Biochem 62, 377–398 (2018).

4. Navas, P. et al. Secondary CoQ10 deficiency, bioenergetics unbalance in disease and aging. Biofactors 47, 551–569 (2021).

5. Wang, Y. & Hekimi, S. The efficacy of coenzyme Q10 treatment in alleviating the symptoms of primary coenzyme Q10 deficiency: A systematic review. J Cell Mol Med 26, 4635–4644 (2022).

6. Luo, J., Yang, H. & Song, B.-L. Mechanisms and regulation of cholesterol homeostasis. Nat. Rev. Mol. Cell Biol. 21, 225–245 (2020).

7. Aronova, S. et al. Regulation of Ceramide Biosynthesis by TOR Complex 2. Cell Metab. 7, 148–158 (2008).

8. Turunen, M., Peters, J. M., Gonzalez, F. J., Schedin, S. & Dallner, G. Influence of peroxisome proliferator-activated receptor α on ubiquinone biosynthesis. J Mol Biol 297, 607–614 (2000).

9. Brea-Calvo, G., Siendones, E., Sánchez-Alcázar, J. A., Cabo, R.de & Navas, P. Cell Survival from Chemotherapy Depends on NF-κB Transcriptional Up-Regulation of Coenzyme Q Biosynthesis. Plos One 4, e5301 (2009).

10. Rensvold, J. W. et al. Defining mitochondrial protein functions through deep multiomic profiling. Nature 606, 382–388 (2022).

11. Lapointe, C. P. et al. Multi-omics Reveal Specific Targets of the RNA-Binding Protein Puf3p and Its Orchestration of Mitochondrial Biogenesis. Cell Syst 6, 125–135 e6 (2018).

12. Veling, M. T. et al. Multi-omic Mitoprotease Profiling Defines a Role for Oct1p in Coenzyme Q Production. Mol Cell 68, 970–977 e11 (2017).

13. Spinazzi, M. et al. PARL deficiency in mouse causes Complex III defects, coenzyme Q depletion, and Leigh-like syndrome. Proc National Acad Sci 116, 277–286 (2019).

14. Ayer, A. et al. Genetic screening reveals phospholipid metabolism as a key regulator of the biosynthesis of the redox-active lipid coenzyme Q. Redox Biol 46, 102127 (2021).

15. Stefely, J. A. et al. Mitochondrial protein functions elucidated by multi-omic mass spectrometry profiling. Nat Biotechnol 34, 1191–1197 (2016).

16. Reidenbach, A. G. et al. Conserved Lipid and Small-Molecule Modulation of COQ8 Reveals Regulation of the Ancient Kinase-like UbiB Family. Cell Chem Biol 25, 154–165 e11 (2018).

17. Lohman, D. C. et al. An Isoprene Lipid-Binding Protein Promotes Eukaryotic Coenzyme Q Biosynthesis. Mol Cell 73, 763–774 e10 (2019).

18. Kuroda, T. et al. FMP30 is required for the maintenance of a normal cardiolipin level and mitochondrial morphology in the absence of mitochondrial phosphatidylethanolamine synthesis. Mol. Microbiol. 80, 248–265 (2011).

19. Ball, W. B. et al. Ethanolamine ameliorates mitochondrial dysfunction in cardiolipin-deficient yeast cells. J. Biol. Chem. 293, 10870–10883 (2018).

20. He, C. H., Xie, L. X., Allan, C. M., Tran, U. C. & Clarke, C. F. Coenzyme Q supplementation or over-expression of the yeast Coq8 putative kinase stabilizes multi-subunit Coq polypeptide complexes in yeast coq null mutants. Biochim Biophys Acta 1841, 630–44 (2014).

21. Floyd, B. J. et al. Mitochondrial Protein Interaction Mapping Identifies Regulators of Respiratory Chain Function. Mol Cell 63, 621–632 (2016).

22. Subramanian, K. et al. Coenzyme Q biosynthetic proteins assemble in a substrate-dependent manner into domains at ER-mitochondria contacts. J Cell Biol 1, 1353–1369 (2019).

23. Eisenberg-Bord, M. et al. The Endoplasmic Reticulum-Mitochondria Encounter Structure Complex Coordinates Coenzyme Q Biosynthesis. Contact (Thousand Oaks) 2, 2515256418825409 (2019).

24. González, J. M. Visualizing the superfamily of metallo-β-lactamases through sequence similarity network neighborhood connectivity analysis. Heliyon 7, e05867 (2021).

25. Merkel, O., Schmid, P. C., Paltauf, F. & Schmid, H. H. O. Presence and potential signaling function of N-acylethanolamines and their phospholipid precursors in the yeast Saccharomyces cerevisiae. Biochimica Et Biophysica Acta Bba - Mol Cell Biology Lipids 1734, 215–219 (2005).

26. Miyata, N., Goda, N., Matsuo, K., Hoketsu, T. & Kuge, O. Cooperative function of Fmp30, Mdm31, and Mdm32 in Ups1-independent cardiolipin accumulation in the yeast Saccharomyces cerevisiae. Sci. Rep. 7, 16447 (2017).

27. Magotti, P. et al. Structure of human N-acylphosphatidylethanolamine-hydrolyzing phospholipase D: regulation of fatty acid ethanolamide biosynthesis by bile acids. Structure 23, 598–604 (2015).

28. Wang, J. et al. Functional Analysis of the Purified Anandamide-generating Phospholipase D as a Member of the Metallo-β-lactamase Family*. J. Biol. Chem. 281, 12325–12335 (2006).

29. Novales, N. A., Meyer, H., Asraf, Y., Schuldiner, M. & Clarke, C. F. Mitochondrial-ER Contact Sites and Tethers Influence the Biosynthesis and Function of Coenzyme Q. Contact 8, 25152564251316350 (2025).

30. Murley, A. et al. ER-associated mitochondrial division links the distribution of mitochondria and mitochondrial DNA in yeast. eLife 2, e00422 (2013).

31. Friedman, J. R. et al. ER Tubules Mark Sites of Mitochondrial Division. Science 334, 358–362 (2011).

32. Nagashima, S. et al. Golgi-derived PI(4)P-containing vesicles drive late steps of mitochondrial division. Science 367, 1366–1371 (2020).

33. Boutry, M. & Kim, P. K. ORP1L mediated PI(4)P signaling at ER-lysosome-mitochondrion three-way contact contributes to mitochondrial division. Nat. Commun. 12, 5354 (2021).

34. Casler, J. C., Harper, C. S., White, A. J., Anderson, H. L. & Lackner, L. L. Mitochondria–ER–PM contacts regulate mitochondrial division and PI(4)P distribution. J. Cell Biol. 223, e202308144 (2024).

35. Kameoka, S., Adachi, Y., Okamoto, K., Iijima, M. & Sesaki, H. Phosphatidic Acid and Cardiolipin Coordinate Mitochondrial Dynamics. Trends Cell Biol. 28, 67–76 (2018).

36. Adachi, Y. et al. Coincident Phosphatidic Acid Interaction Restrains Drp1 in Mitochondrial Division. Mol. Cell 63, 1034–1043 (2016).

37. Choi, S.-Y. et al. A common lipid links Mfn-mediated mitochondrial fusion and SNARE-regulated exocytosis. Nat. Cell Biol. 8, 1255–1262 (2006).

38. Pettinati, I., Brem, J., Lee, S. Y., McHugh, P. J. & Schofield, C. J. The Chemical Biology of Human Metallo-β-Lactamase Fold Proteins. Trends Biochem. Sci. 41, 338– 355 (2016).

39. Freyer, C. et al. Rescue of primary ubiquinone deficiency due to a novelCOQ7defect using 2,4–dihydroxybensoic acid. Journal of Medical Genetics 52, 779–783 (2015).

40. Murray, N. H. et al. 2-Propylphenol Allosterically Modulates COQ8A to Enhance ATPase Activity. Acs Chem Biol 17, 2031–2038 (2022).

41. Castellani, B. et al. Synthesis and characterization of the first inhibitor of N-acylphosphatidylethanolamine phospholipase D (NAPE-PLD). Chem Commun (Camb) 53, 12814–12817 (2017).

42. Longtine, M. S. et al. Additional modules for versatile and economical PCR-based gene deletion and modification in Saccharomyces cerevisiae. Yeast 14, 953–961 (1998).

43. Gietz, R. D. & Schiestl, R. H. High-efficiency yeast transformation using the LiAc/SS carrier DNA/PEG method. Nat. Protoc. 2, 31–34 (2007).

44. Meisinger, C., Pfanner, N. & Truscott, K. N. Isolation of Yeast Mitochondria. Methods in Molecular Biology 313, 33–39 (2006).

45. Kemmerer, Z. A. et al. UbiB proteins regulate cellular CoQ distribution in Saccharomyces cerevisiae. Nat Commun 12, 4769 (2021).

46. Tyanova, S. et al. The Perseus computational platform for comprehensive analysis of (prote)omics data. Nat Methods 13, 731–740 (2016).

47. Hutchins, P. D., Russell, J. D. & Coon, J. J. LipiDex: An Integrated Software Package for High-Confidence Lipid Identification. Cell Syst. 6, 621-625.e5 (2018).

48. Hutchins, P. D., Russell, J. D. & Coon, J. J. Mapping Lipid Fragmentation for Tailored Mass Spectral Libraries. J. Am. Soc. Mass Spectrom. 30, 659–668 (2019).

49. Anderson, B. J., Brademan, D. R., He, Y., Overmyer, K. A. & Coon, J. J. LipiDex 2 Integrates MSn Tree-Based Fragmentation Methods and Quality Control Modules to Improve Discovery Lipidomics. Anal. Chem. 96, 6715–6723 (2024).

50. Guo, X. et al. Ptc7p Dephosphorylates Select Mitochondrial Proteins to Enhance Metabolic Function. Cell Rep 18, 307–313 (2017).

